# Social affective behaviors among female rats involve the basolateral amygdala and insular cortex

**DOI:** 10.1101/2023.02.02.526780

**Authors:** Anthony Djerdjaj, Nathaniel S. Rieger, Bridget H. Brady, Bridget N. Carey, Alexandra J. Ng., John P. Christianson

**Affiliations:** Department of Psychology & Neuroscience, Boston College, Chestnut Hill, MA 02467

**Keywords:** empathy, stress, social, rat, neural circuit, oxytocin

## Abstract

The ability to detect, appraise, and respond to another’s emotional state is essential to social affective behavior. This is mediated by a network of brain regions responsible for integrating external cues with internal states to orchestrate situationally appropriate behavioral responses. The basolateral amygdala (BLA) and the insular cortex are reciprocally connected regions involved in social cognition and prior work in male rats revealed their contributions to social affective behavior. We investigated the functional role of these regions in female rats in a social affective preference (SAP) test in which experimental rats approach stressed juvenile but avoid stressed adult conspecifics. In separate experiments, the BLA or the insula were inhibited by local infusion of muscimol (100ng/side in 0.5μL saline) or vehicle prior to SAP tests. In both regions, muscimol interfered with preference for the stressed juvenile and naive adult, indicating that these regions are necessary for appropriate social affective behavior. In male rats, SAP behavior requires insular oxytocin but there are noteworthy sex differences in the oxytocin receptor distribution in rats. Oxytocin (500nM) administered to the insula did not alter social behavior but oxytocin infusions to the BLA increased social interaction. In sum, female rats appear to use the same BLA and insula regions for social affective behavior but sex differences exist in contribution of oxytocin in the insula.

## Introduction

The capacity of one individual to detect and make decisions based on the emotional or arousal state of another is key to understanding both healthy and pathological social interactions. In humans, these processes are thought to be fundamental to empathic cognition. When describing such elementary parts of empathy, de Waal argued, “The lowest common denominator of all empathic processes is that one party is *affected* by another’s emotional or arousal state” (de Waal, 2008, p.282). This primal, automatic process is sensory in nature and it, by definition, precedes the higher-order manifesta­tions of empathy that are widely studied in humans (2) and increasingly observed in laboratory rodents (3–7). Thus, to detect the emotion of another is the precursor to emotion contagion, emotion recognition, empathic helping, perspective taking, and so on, all of which shape an individual’s reactions to and interactions with others. Abnormalities in these ‘higher’ processes are central to numerous psychiatric conditions including autism spectrum disorder (ASD), Fragile X, depression, psychopathy, schizophrenia, borderline personality disorder and alexithymia (8–12). Relevant to the present study, there are marked sex differences in prevalence of social dysfunction in psychopathology (13–16) making the translational study into sex differences in the neurobiology of social affective behaviors a high priority.

To investigate the neurobiology of simple empathy-like processes in a translational model, our laboratory developed a social affective preference (SAP) test (17–20). The SAP test allows for the observation of a rat’s unconditioned response to a pair of stimuli rats, one of which had been exposed to a stressor and the other naïve to treatment. Interestingly, when given the choice between naïve and stressed juveniles, adult test rats spend more time exploring the stressed juveniles, but this preference was exactly the opposite if the stimuli rats were post pubertal adults. We conducted extensive behavioral ethography of the experimental and conspecific rat behaviors including ultrasonic vocalizations (17); and concluded that the approach behavior of the experimental rat is a reliable dependent measure that is influenced by the social affective signals of the conspecifics. Thus, the SAP test reflects something well established with regard to human social behavior: features of the target stimulus, including age, are critical determinants to whether or not an individual will approach, help or avoid another in distress (21).

In our first behavioral studies, we included both male and female rats and the pattern of approach to stressed juveniles and avoidance of stressed adults was present in both sexes within the same testing parameters (17,22). However, when we began to investigate neurobiological mechanisms, which entail more invasive and stressful procedures such as cannula implant surgeries and injections, female SAP behavior became unreliable while male behavior was robust. This led us to pursue neural mechanisms first in males (reviewed below) while seeking parameters for female SAP tests that would be compatible with modern neuroscience technologies.

Our studies in male rats initially focused on the insula. The insula is central to the prevailing neuroanatomical models of empathy, which have substantial support from human neuroimaging studies (4,23). Classically identified as a neural locus of somatosensory interoception ((Bud) Craig & D, 2009; W. G. Chen et al., 2021), there has been an explosion of fMRI studies over the last 10+ years implicating insula in an impressive range of cognitive processes (26), including but not limited to emotion recognition, salience detection, pain, drug craving and so on (27–31). It is likely that many of these can be tied to an elementary function of integration and salience detection, a consequence of insula’s multisensory inputs and reciprocal connectivity with brain networks serving attention, arousal, action, and emotion (32). Thus, we hypothesized that behavior towards or away from stressed conspecifics in the SAP test would require the insula. Indeed, interaction with stressed conspecifics increases activity in the insula and pharmacological, optogenetic, and chemogenetic inhibition all abolish preference behaviors (17– 20,33). Importantly, oxytocin and corticotropin releasing factor, peptides implicated in social behavior (34) and stress responses (35), respectively, were found to be necessary modulators of insular synaptic physiology and the corresponding receptors necessary for social affective preference behaviors (17,33). Subsequently, numerous other reports have shown important roles for the insula in social affective behaviors (36–38) in males.

We interpret the importance of the insula to social affective behaviors as a consequence of its interconnections with the network of highly conserved brain regions that orchestrate social behavior (39) and our recent work has addressed the necessity of this connectivity. The basolateral amygdala (BLA) is a key component of several network models of social cognition including the Social Decision Making Network (40) and “social brain” (41) and sends a dense fiber input to the posterior insula (42,43). BLA neurons are remarkably sensitive to conspecific behavior (44); respond during direct observation of a conspecific receiving shock (45); and are activated upon exposure to a stressed conspecific (46,47). We hypothesized that detection or appraisal of social stress stimuli and the integration of this external valence information with internal states would require the BLA and its projections to the insula, respectively. In males, chemogenetic inhibition of the BLA and its projection terminals in the insula did in fact abolish social preference for stressed juveniles (20), providing a mechanistic basis for the detection, appraisal, and behavioral response to emotionally salient stimuli.

Sex differences in the prevalence and manifestation of neuropsychiatric disorders such as ASD and schizophrenia (15,16,48–51) warrant research into the corresponding neurobiology of socioemotional behavior in females. Underlying differences in functional connectivity during emotion recognition and empathy-related processes are thought to contribute to sex-specific manifestations of psychiatric disorders (52,53). For example, amygdala-insula connectivity is increased in human females when viewing the faces of stressed others compared to males (54). Sex differences in network function could be attributed to sex differences in the distribution of oxytocin fibers and receptors. Male rats have greater oxytocin binding in the agranular insula, indicating a potential sex-specific role of oxytocin in the insula (55). Less is known about differences in oxytocin receptor expression and function in the BLA, though other amygdala subnuclei, such as the medial and central amygdala, display sex-specific oxytocin binding densities and functions (55–57). Insular oxytocin in males increases excitatory synaptic transmission and is necessary for appropriate social affective preference behavior, while its role in female insula remains unknown (17).

The goal of the current study was to determine if female rats utilize the same neural systems to navigate social affective decision making that we have discovered in males. As noted above, prior efforts to study these systems in females failed because the behavior phenotype was not robust to surgical procedures and injections. To overcome this we lengthened the time allotted for recovery from surgery to 4 weeks and acclimated the rats to handling and experiment interaction for 15 days prior to SAP tests (58). Here, we infused muscimol, the GABA_A_ receptor agonist, to the insula or BLA of female rats for reversible pharmacological inhibition during SAP testing with either juvenile or adult isosexual conspecifics. We also sought to ascertain whether oxytocin affects females similarly to males by infusing oxytocin into the insula or BLA prior to one-on-one social interaction tests.

## Materials and Methods

### Animals

All rats used were female Sprague-Dawleys purchased from Charles River Laboratories (Wilmington, MA). All rats were allowed to acclimate to the vivarium in the Boston College Animal Facility for at least 7 days prior to any procedure being carried out. Experimental subjects arrived weighing 225-250 grams and were housed in pairs. Juvenile conspecifics arrived at PN21 and adult conspecifics arrived at PN55. Conspecifics were housed in groups of 3. Food and water were available *ad libitum*. Rats were housed on a 12h light/dark cycle with behavioral experiments occurring within the first 4 h of the light cycle. All procedures were approved by the Boston College Institution Animal Care and Use Committee and adhered to the Public Health Service *Guide for the Care and Use of Laboratory Animals*.

### Surgical Procedures - Cannula Implantation

To inhibit insula and BLA activity indwelling cannula were implanted bilaterally into either the insula or BLA to allow for direct infusion of muscimol or vehicle into each region. Experimental adult females underwent surgery under inhaled anesthesia (2-5% v/v isoflurane in O_2_). Bilateral cannula (Plastics One) were implanted into the posterior insula (from bregma: A/P −1.8, M/L +/−6.5, D/V −6.9) or the BLA (from bregma: A/P −2.2, M/L +/− 4.5, D/V −8.4). Cannula were fixed in place with stainless steel screws and acrylic cement and were fitted with stylets to maintain patency. After surgery, rats were subcutaneously injected with meloxicam (1 mg/kg, Eloxiject; Henry Schein) as well as the antibiotic penicillin (1 mg/kg, Combi-Pen; Henry Schein). 10 mL lactated Ringer’s solution was delivered in two doses of 5 mL subcutaneously on the right and left side of the body. Rats were allowed 4 weeks of recovery before undergoing behavioral procedures as described below. 3 weeks prior to behavioral testing, test rats were wrapped in a towel and handled by an experimenter for 1 minute each weekday for a total of 15 days of handling. This was done to ensure rats were fully habituated to handling prior to drug infusion. To inhibit either the insula or the BLA, muscimol (100 ng/side in 0.5mL of saline) was microinjected 1 hour prior to testing. We have used this dose previously (17,59,60).

### Social Affective Preference (SAP) Test

This procedure allows for the observation of a rodent’s behavior when presented with conspecifics of varying social affect. The SAP paradigm takes place in a plastic arena (76.2cm × 20.3cm × 17.8 cm, Length × Width × Height) with beta chip bedding and a transparent plastic lid. Conspecifics were individually placed into clear acrylic chambers (18 × 21 × 10cm; L × W × H) comprised of acrylic rods spaced 1 cm apart to allow for direct interaction. These chambers are placed at opposite ends of the plastic arena during testing. Testing consists of 2 habituation days followed by 2 test days. On day 1, experimental rats were placed in the testing room for 1 hr and exposed to the testing arena for 15 minutes before being returned to their home cage. On day 2, experimental rats were placed in the plastic testing arena for 5 minutes of behavior testing where they were presented with two naïve juvenile or adult conspecifics in the acrylic chambers. Days 3 and 5 test the experimental rats’ social affective preference after injection of vehicle or muscimol, as described above. Rats were placed in the arena for 1 h and then presented with a pair of unfamiliar conspecifics. One conspecific received an acute stressor of 2 footshocks immediately preceding placement in its chamber of the testing arena (5s, 1 mA, inter-shock interval of 50s); the other conspecific was naive to any treatment. A trained observer quantified the amount of time the experimental rat spends investigating each conspecific. Social investigation was defined as time spent sniffing or touching the conspecific through the acrylic bars. After tests on day 3, rats were returned to their home cage and left undisturbed for 48 h to ensure muscimol cleared. Testing on day 5 was the same with rats receiving the opposite drug treatment as on day 3 in a counterbalanced, within-subjects design. All tests were recorded in digital video. Recordings were scored by a trained observer blind to the experimental conditions to establish inter-rater reliability.

### One-on-one social interaction tests

Three days after SAP testing, the same experimental rats underwent one-on-one social exploration tests. Each experimental rat was placed into a standard plastic tub cage with beta chip bedding and a wire lid 1 h prior to testing. Testing consisted of a naive juvenile or adult being introduced into the experimental rat’s cage for 5 minutes. Exploratory behaviors (sniffing, pinning, allogrooming) initiated by the experimental rat were timed by an observer. Each experimental rat was given tests on consecutive days, once 15 minutes after receiving bilateral infusions of oxytocin (0.5 μL of 500 nM in 0.9% saline vehicle equivalent to 250 pg oxytocin per side) and once after receiving bilateral infusions of the vehicle. This time and dose was selected based on our prior studies (17). Drug injection order was counterbalanced.

### Cannula placement verification

After behavioral testing was finished, experimental rats were overdosed with tribromoethanol and decapitated. Directly prior to this, a vaginal lavage was taken from each rat to identify the day of estrous. After decapitation, the brains were removed, and flash frozen for slicing. Brains were sectioned at 40 μm using a freezing cryostat (Leica CM1860 UV) and slices were mounted on gelatin-coated slides. A cresyl violet stain was performed for verification of cannula placement under a microscope.

### Statistical analysis

Social interactions were defined as sniffing or touching of the conspecifics and timed by experimenters blind to treatment. Inter-rater reliability was regularly established. Data from experimental rats were only included if site-specificity criteria were met after verification of cannula placement. For insula cannula, data were included if the lowest point of cannula damage was found in the posterior insula. For BLA injections, data were included if the lowest point of cannula damage was found in the posterior BLA. It is important to note that some rats only had unilateral injections but were included if cannula placement was correct in order to minimize the amount of animals used. Social interaction and preference behaviors were analyzed using a repeated measures Analysis of Variance (ANOVA). Main effects and interaction effects were deemed significant at p < 0.05 and followed by Sidak post-hoc tests to maintain experiment-wise type 1 error rate to ɑ < 0.05. Preference for the stressed conspecific in each condition was calculated as a percentage of the total time spent investigating both conspecifics and compared with ANOVA. One-on-one social interaction data was analyzed using paired t-tests. Preference for social interaction under oxytocin was calculated as a percentage of time spent interacting under vehicle and analyzed using one sample t-tests. Statistical analyses were conducted with Prism 9 (Graphpad Software).

## Results

### Insula is necessary for social approach to stressed juveniles and naïve adults in female rats

To determine the effect of insula inhibition on female social affective preference for juveniles, adult female experimental rats received bilateral cannula implants in the insula and later underwent SAP testing with female juvenile conspecifics after vehicle or muscimol injections (Fig 1A). After cannula verifications, 12 rats met the inclusion criteria (Fig 1B). The amount of time the experimental rat spent investigating the naïve and stressed conspecific was analyzed with ANOVA where drug treatment (vehicle vs. muscimol) and conspecific affect (naïve vs. stressed) were treated as within-subjects factors. Experimental rats preferred interaction with stressed juveniles after vehicle injection but appeared to lose this preference after muscimol administration (Fig 1). There were main effects of both drug treatment (*F*(1,11) = 4.90, p = 0.049, ⌷^2^ = 6.57) and social affect (*F*(1,11) = 9.33, p = 0.011, ⌷^2^ = 16.5) as well as a drug by affect interaction (*F*(1,11) = 7.42, p = 0.020, ⌷^2^ = 6.77). Post-hoc comparison revealed a significant difference between social investigation of naive and stressed juveniles in the vehicle condition (p = 0.0009) that was not present in the muscimol condition (p = 0.517, Fig 1C). A separate cohort of adult females with insula cannula were tested with adult conspecifics. After cannula verifications, 9 rats met inclusion criteria (Fig 1B) and were analyzed as above resulting in a significant drug by affect interaction (*F*(1,8) = 98.3, p < 0.0001, ⌷^2^ = 30.5). Post-hoc comparisons revealed that experimental rats spent significantly more time investigating naïve adults in the vehicle condition (p < 0.0001) and significantly less time investigating naïve adults in the muscimol condition (p = 0.006, Fig 1D).

**Figure 1.**
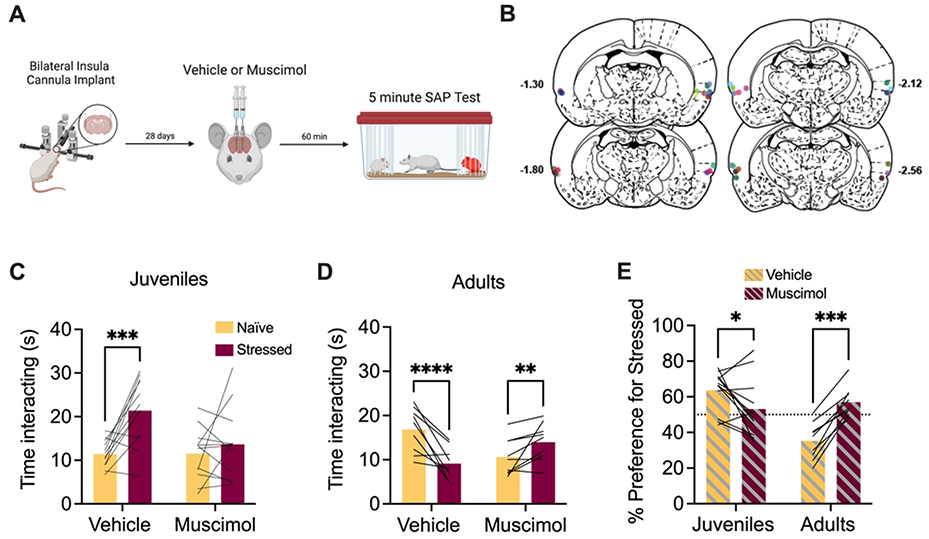
Pharmacological inactivation of the insula in female rats interfered with social affective preference behavior. **A**. Schematic diagram of experimental design. Bilateral cannula were implanted into the insula of female test rats and, 4 weeks later, rats received saline or muscimol infusions 60 minutes before SAP testing with juvenile or adult conspecifics. **B**. Schematic of cannula implants into the posterior insula. **C**. Mean (with individual replicates) time spent interacting with the naïve or stressed juvenile conspecifics during the 5 min. SAP test. After vehicle injections, rats preferred interactions with stressed juveniles compared to naïve juveniles (p = 0.0009), which was abolished after muscimol infusion (100 ng per side in 0.5μL saline). **D**. Mean (with individual replicates) time spent interacting with the naïve or stressed adult conspecifics during the 5 min SAP test. After vehicle injections, rats preferred interactions with naïve adults compared to stressed adults (p < 0.0001), which was reversed by muscimol infusions (p = 0.006). **E**. Data from C and D presented as a mean (with individual replicates) preference for the stressed conspecific as a percentage of total social interaction to allow a comparison of drug effect by age. Experimental rats preferred interactions with stressed juveniles (p = 0.028) and naïve adults (p = 0.0002) under vehicle treatment, which was abolished following muscimol infusion. *p < 0.05, **p < 0.01, ***p < 0.001, ****p < 0.0001. Diagram in A created with BioRender.com. Atlas images recreated from Paxinos & Watson (1998).

The preference for the stressed conspecific in each experiment was calculated as a percentage of the total time spent investigating both conspecifics. A 2-way ANOVA was performed with drug treatment (vehicle vs. muscimol) as a within-subjects variable and age (juvenile vs. adult) as a between-subjects variable. This revealed a main effect of age (*F*(1,19) = 7.25, p = 0.014, ⌷^2^ = 15.3) and a drug by age interaction effect (*F*(1,19) = 29.7, p < 0.0001, ⌷^2^ = 26.5). Post-hoc comparisons revealed a significant change in preference for stressed juveniles (p = 0.028) and naïve adults (p = 0.002) when comparing vehicle to muscimol (Fig 1E). In summary, insula muscimol infusions in female rats interfered with social preference for stressed juveniles and naïve adults.

### BLA inhibition abolishes social preference for stressed juveniles and naïve adults in female rats

To determine the effects of BLA inhibition on social approach to stressed juveniles, bilateral cannula were implanted into the BLA of adult female rats through which muscimol was injected 1 h prior to testing (Fig 2A). Twelve rats met inclusion criteria after cannula verifications (Fig 2B). It is important to note that, to minimize the number of test subjects used, 3 rats with correct unilateral cannula placements were included. Similar to above, experimental rats preferred interaction with stressed juveniles after vehicle injection but appeared to lose this preference after muscimol administration (Fig 2C). The amount of time spent investigating the naive and stressed conspecifics was recorded and a 2-way ANOVA with drug treatment (vehicle vs. muscimol) and social affect (naive vs. stressed) as within-subjects variables resulted in a significant drug by social affect interaction (*F*(1,11) = 7.04, p = 0.022, ⌷^2^ = 9.00). Post-hoc comparisons did not yield any significant differences, although the difference in time spent investigating the stressed vs. the naive juvenile in the vehicle condition approached significance (p = 0.054, Fig 2C).

**Figure 2.**
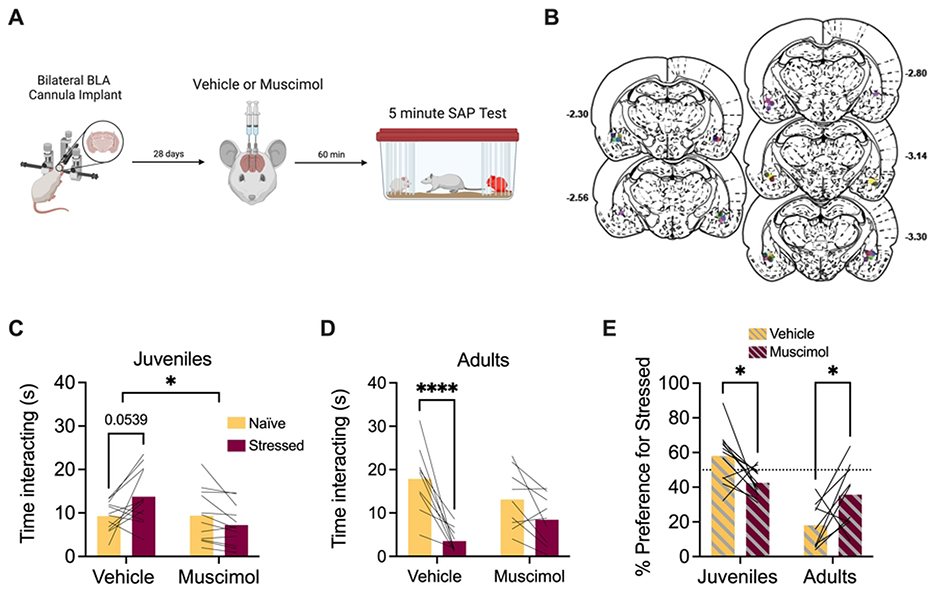
Pharmacological inactivation of the BLA in female rats interfered with social affective preference behavior. **A**. Schematic diagram of experimental design. Bilateral cannula were implanted into the BLA of female test rats and, 4 weeks later, rats received saline or muscimol infusions 60 minutes before SAP testing with juvenile or adult conspecifics. **B**. Schematic of cannula implants into the BLA. **C**. Mean (with individual replicates) time spent interacting with the naïve or stressed juvenile conspecifics during the 5 min. SAP test. After vehicle injections, rats preferred interactions with stressed juveniles compared to naïve juveniles (p = 0.054), which was abolished after muscimol infusion (100 ng per side in 0.5μL saline) A social affect x drug interaction effect was observed (p = 0.022). **D**. Mean (with individual replicates) time spent interacting with the naïve or stressed adult conspecifics during the 5 min. SAP test. After vehicle injections, rats preferred interactions with naïve adults compared to stressed adults (p < 0.0001), which was reduced following muscimol infusion. **E**. Data from C and D presented as a preference for the stressed conspecific as a percentage of total social interaction (with individual replicates). Experimental rats preferred interactions with stressed juveniles (p = 0.038) and naïve adults (p = 0.04) under vehicle treatment, which was abolished following muscimol infusion. *p < 0.05, ****p < 0.0001. Diagram in A created with BioRender.com. Atlas images recreated from Paxinos and Watson (1998).

Notably, in this experiment 4 of 12 females avoided the stressed juvenile in the vehicle condition which is a slightly larger portion than typically observed but consistent with our prior work where preference fell along a normal distribution with some animals avoiding juveniles (17). A separate cohort of adult female rats were tested with adult conspecifics and analyzed as above; 9 rats, including 4 with correct unilateral cannula placements, met inclusion criteria after cannula verifications (Fig 2B). The ANOVA resulted in a significant main effect of social affect (*F*(1,8) = 18.95, p = 0.0024, ⌷^2^ = 37.0) and a drug by social affect interaction effect (*F*(1,8) = 14.04, p = 0.0057, ⌷^2^ = 9.52). Post-hoc comparisons revealed that experimental rats spent significantly more time investigating naïve adults in the saline condition (p < 0.0001) (Fig 2D).

The preference for the stressed conspecific was calculated as described above and a 2-way ANOVA with drug treatment (saline vs. muscimol) as a within-subjects variable and age (juvenile vs. adult) as a between-subjects variable resulted in a main effect of age (*F*(1,19) = 34.9, *p* < 0.0001, ⌷^2^ = 35.9) and a drug by age interaction effect (*F*(1,19) =12.9, p=0.0019, ⌷^2^ = 18.0). Post-hoc comparisons revealed a significant change in preference for stressed juveniles (p = 0.038) or naïve adults (p = 0.04) between vehicle and muscimol (Fig 2E). In all, these results indicate that BLA infusions of muscimol impair approach to stressed juveniles and avoidance of stressed adults in female rats.

### Oxytocin in the BLA, but not the insula, increases social interaction with naïve juvenile and adult conspecifics

The foregoing results begin to establish that female rats use some of the same brain regions found to be important for social affective behaviors in males. In our first investigations of this neurobiology in male rats, we found that oxytocin was a necessary and sufficient modulator of the insular cortex. Given known sex differences in oxytocin receptor expression (55,57,62), we next sought to test whether oxytocin would alter social interactions in females. To determine whether oxytocin infusions to the insula or BLA effect social behavior in female rats, 3 days after SAP testing, 10 rats with bilateral insula cannula and 21 rats with BLA cannula received infusion of either vehicle or oxytocin (500 nM) 15 minutes prior to one-on-one social interaction tests (Fig 3A). Initially these tests were conducted with juvenile conspecifics (n=10 for insula; n=12 for BLA; see Figs 1B and 2B for cannula verifications). Because oxytocin in the BLA augmented social interaction with juveniles we then repeated the experiment for BLA oxytocin infusion with adult conspecifics (n = 9; see Fig 2B for cannula verifications). Time spent exploring the naïve conspecific over the course of the 5 min test was recorded. Initially, a 2-way ANOVA with drug treatment (vehicle vs. oxytocin) as a within-subjects variable and age (juvenile versus adult) as a between-subjects variable was performed on the data obtained from rats with BLA cannula, revealing a main effect of drug (*F*(1,19) =12.8, p = 0.002, ⌷^2^ = 18.9) but no significant effect of conspecific age or interactions. Therefore, data from BLA juvenile and adult social interaction tests were pooled together to compare the effect of oxytocin across regions. In this case an ANOVA with drug treatment (vehicle vs. oxytocin) as a within-subjects variable and brain region (insula versus BLA) as a between-subjects variable resulted in main effects of drug (*F*(1,29) = 8.29, p = 0.007, ⌷^2^ = 0.04) and brain region (*F*(1,29) = 13.9, p = 0.0008, ⌷^2^ = 0.25). Post-hoc comparisons revealed a significant increase in social interaction after oxytocin infusion into the BLA compared to vehicle (p = 0.0008) with no difference in the insula (p = 0.72, Fig 3B). Time spent exploring after oxytocin was converted to a percentage of vehicle exploration time and one sample t-tests were conducted comparing the mean percentage of time spent exploring under oxytocin to a theoretical mean of 100%. Social interaction did not increase after infusion of oxytocin into the insula but did increase after infusion into the BLA (p = 0.001, Fig 3C).

**Figure 3.**
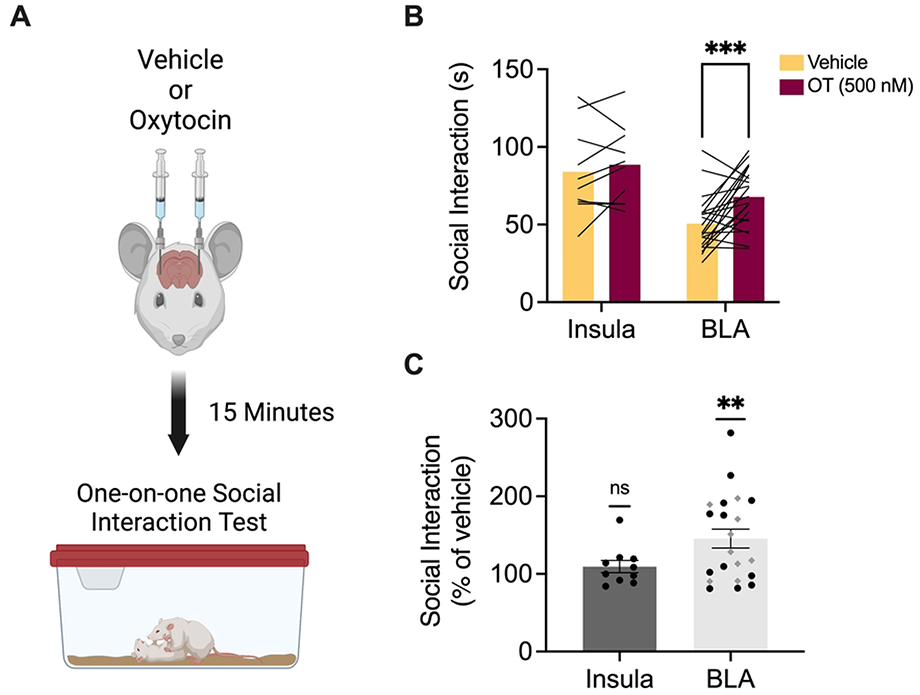
Oxytocin infusion into the BLA, but not the insula, of female test rats increased social interaction with naïve juvenile and adult conspecifics. **A**. Schematic diagram of experimental design. One week after SAP testing, female test rats with bilateral cannula implants in either the insula or BLA received saline or oxytocin (500 nM) infusions 15 min prior to one-on-one social interaction tests with naïve juvenile or adult conspecifics. **B**. Mean (with individual replicates) time spent interacting with naïve juvenile and adult conspecifics during a 5 min social interaction test after vehicle or oxytocin infusion into the insula or BLA. Oxytocin in the insula had no effect on social interaction with juvenile conspecifics compared to social interaction after vehicle infusions. Oxytocin in the BLA significantly increased social interaction with both juvenile and adult conspecifics (p = 0.0008). **C**. Data from panel B presented as social interaction under oxytocin as a percentage of social interaction under vehicle (with SEM). Oxytocin in the BLA produced a significant increase in social interaction with juveniles (black circles) and adults (gray diamonds) compared to 100% (p = 0.0013). **p < 0.01, ***p < 0.001. Diagram in A created with BioRender.com.

Oxytocin infusions to the BLA had noticeably variable effects which may be attributable to fluctuations in oxytocin receptor expression associated with estrus phase (55). We analyzed the BLA data again with subjects grouped by estrus phase. An ANOVA with drug (vehicle vs. oxytocin) as a within-subjects variable and estrus phase (proestrus vs. diestrus vs. metestrus vs. estrus) as a between-subjects variable revealed a main effect of drug (*F*(1,17) = 7.4, *p* = 0.014, ⌷^2^ = 11.8) but no significant effect of phase or drug by phase interactions (data not shown). Thus, estrus phase does not appear to mediate the variability in oxytocin efficacy on social interaction when infused to the BLA.

## Discussion

We investigated whether posterior insula or BLA inhibition in female rats interfered with preference for stressed juveniles and naïve adults typically observed in a social affective preference test (17,20). Akin to similar results in males, muscimol infusions into either the insula or the BLA reduced time spent interacting with stressed juveniles and naïve adults, respectively. In contrast to its prosocial effects in males, infusion of oxytocin into the insula had no effect on social interaction with juvenile conspecifics in females, suggesting a sex-specific role of this neuropeptide in social behavior. Alternatively, oxytocin infusion into the BLA of female rats increased social interaction with both juvenile and adult conspecifics, establishing a potential prosocial role of BLA oxytocin in females. These results add to our understanding of the neural circuit basis for social emotional behavior in rats.

The decision to approach or avoid a conspecific is informed by internal signals present within the observer and external cues emitted from the target conspecific. Interoception and salience detection are mediated in part by the insula (32), an accessory node to the social decision-making network (39). Anatomical studies incorporating both male and female mice have reported no major sex differences in insula connectivity with other brain regions (63), supporting the idea that similar anatomical structure may give rise to similar functions across sexes. Indeed, we’ve shown here that posterior insula activity in adult females is necessary for social affective preference for stressed juveniles and naïve adults, like our findings in male rats (17). Given its position as an intersection for multisensory information processing, it’s likely that the insula guides social affective behavior through its integration of external stimuli and internal states.

External cues such as ultrasonic vocalizations (64,65), chemosignals (66), and overt behaviors convey a conspecific’s affective state and influence an observer’s reactions. Vocalizations are especially important to social interaction as they signal positive or negative affect to observers. In rats, 22-Hz calls are associated with aversive states while 50-Hz calls signal positive states (67). In males, the BLA responds differentially to each call, increasing activity to 22-Hz calls and decreasing activity to 50-Hz calls (68). While BLA responses to vocalizations in females are unknown, female rodents convey affective states via vocalizations in social situations similar to males (69–71). Chemosignals are also essential to social behavior and BLA inhibition in females abolishes prioritization of social odorants over nonsocial odorants, specifically urine from female conspecifics (72). Given its known role as a valence detector in males, it’s likely that the BLA acts similarly in females, encoding positive or negative affective states of others based on socially derived cues.

Social stimuli can be appraised as either potentially rewarding (*e.g*., a mate) or hazardous (*e.g*., a dominant male) which would likely lead to opposite social behaviors, approach or avoidance, respectively. In the SAP test, juvenile stress signals may be perceived as non-threatening and motivate parental stress-buffering approach behaviors, while adult stress signals might be perceived as cues of imminent social danger and motivate defensive or avoidant behaviors. In a prior analysis of Fos in male rats that underwent social interactions with either naive or stressed juveniles or adults, we observed the correlation between BLA and insular Fos to vary based on the age and stress of the conspecific (17) and inhibition of BLA synaptic terminals within the posterior insula interfered with social approach (20) suggesting that this is a functionally important circuit and insula-projecting amygdala neurons relay information about the emotional value of the social stimuli to the insular cortex.

An inference made in our prior work is that engagement with stressed conspecifics evokes a specific pattern of activity within the insula and the associated social decision-making network that is distinct from the pattern of activation that results from interactions with naive conspecifics. In males, neuromodulators that are well established contributors to social and stress-related behaviors, oxytocin and corticotropin releasing factor, augment insular cortex intrinsic and synaptic excitability and their receptors are necessary for typical social affective preference (17,58). Sex differences in the oxytocin system are common (56,73) and there are age (55,57,62) and maternal (74) factors that affect cortical oxytocin receptor functions. In Wistar rats, adult males have greater oxytocin receptor binding compared to females (Dumais et al., 2013, but see also Smith et al., 2017) providing a possible explanation for a lack of effect of oxytocin infusions on social interaction here. We also investigated the effect of oxytocin on synaptic transmission in acute insular cortex slices. In males, 500 nM oxytocin causes a robust increase in synaptic efficacy as measured by field excitatory postsynaptic potentials, but neither 500 nM or 2 μM doses had effects on female insula slices (B. Brady & J. Christianson unpublished data). The current data suggest that oxytocin is not important to insula function in female rats. Interestingly, oxytocin had prosocial effects when administered to the BLA of females. A hypothesis to test in future studies would be that oxytocin released during interactions with stressed juveniles augments BLA neurons projecting to the insula providing a mechanism to drive insula activity. Looking beyond the oxytocin system, neuromodulators that define the social decision-making networks, including vasopressin, opioids, and dopamine (75,76), should be explored in the female brain.

The SAP phenomenon tested here rests on the finding that experimental rats prefer interactions with stressed over naïve juveniles and naïve over stressed adults (17). While the exact impetus for this behavior in rodents is unknown, work done in both humans and rodents grants us insight into potential motivations and sex differences for this social decision-making process. Human neuroimaging in parents revealed correlated activity in a network of brain regions that mediate parental impulses and behaviors (77), though women display a greater incentive salience toward infants (78) and exhibit stronger amygdala activation in response to infant vocalizations than men (79). In rodents, virgin females are more prone to parental behaviors than virgin males and fluctuations in hormones following gestation increases maternal behaviors in females (80,81). The BLA may be a crucial mediator of this behavior, as lesions impaired maternal behaviors toward pups in virgin female mice (82). Taking these findings into consideration, it is likely that prosocial behavior towards stressed juveniles could be motivated by parental responsiveness, and this may be more pronounced in females. On the other hand, adults may avoid stressed adults as a defensive behavior, serving to avoid close contact with aggressive or otherwise harmful others. Neither the current data or our prior work with males has uncovered the neural circuits that mediate avoidance of adult conspecifics in the SAP test beyond the insula. Sex differences exist in the circuitry governing social avoidance behaviors and oxytocin acts as a sex-specific modulator of these behaviors (83). Therefore, actions of oxytocin elsewhere in the social decision-making network could contribute to avoidance, providing a starting point to expand the neural circuit mechanism of social affective decision-making.

## Acknowledgements

The authors wish to thank Natalie Cortopassi who made important contributions to the pilot studies and to Nancy McGilloway and Todd Gaines who provided excellent animal husbandry in the Boston College Animal Care Facility. Funding in support of this work came from National Institute of Mental Health grant MH119422. Bridget Brady and Bridget Carey were supported by Boston College Undergraduate Research Fellowships.

